# Does extracellular DNA mask microbial responses to a pulse disturbance?

**DOI:** 10.1101/2021.12.04.471228

**Authors:** HA Kittredge, KM Dougherty, K Glanville, SE Evans

**Affiliations:** W.K. Kellogg Biological Station, Department of Integrative Biology, Michigan State University, Hickory Corners, MI 49060; Program in Ecology, Evolution, and Behavior, Michigan State University, East Lansing, MI 48824; W.K. Kellogg Biological Station, Department of Plant, Soil, and Microbial Sciences, Michigan State University, Hickory Corners, MI 49060; Microbiology and Molecular Genetics, Michigan State University, East Lansing, MI 48824

**Keywords:** extracellular DNA (exDNA), soil microbial communities, drying-rewetting, pulse disturbance, necromass

## Abstract

A major goal in microbial ecology is to predict how microbial communities will respond to global change. However, DNA-based sequencing that is intended to characterize live microbial communities includes extracellular DNA (exDNA) from non-viable cells. This could obscure relevant microbial responses, particularly to pulse disturbances which kill bacteria and have disproportionate effects on ecosystems. Here, we characterize bacterial communities before and after a drying-rewetting pulse disturbance, using an improved method for exDNA exclusion. We find that exDNA removal is important for detecting subtle yet significant changes in microbial abundance, diversity, and community composition across the disturbance. However, inclusion of exDNA did not obscure results to a large extent, only sometimes altering statistical significance but rarely changing the direction of the response or general conclusions about bacterial disturbance dynamics. Although there may be instances where exDNA removal is essential for accurate representation of microbial communities, our study suggests these scenarios will be difficult to predict *a priori*. Overall, we found no evidence that certain time points across the distrubance were more affected by exDNA inclusion, nor did the size or composition of exDNA pools accurately predict when exDNA would alter significance levels. However, exDNA dynamics did vary strongly across the two soil types tested.

## Introduction

Large pools of prokaryotic extracellular DNA (exDNA) can accumulate in the environment as bacteria die (1–5). This exDNA is included in molecular characterizations of ‘live’ communities, potentially altering estimates of bacterial abundance, diversity, and community composition (6–11). Despite this danger, there are surprisingly few assessments of exDNA and live DNA, which are needed to understand when and where inclusion of exDNA can obscure our understanding of microbial dynamics. Our understanding of bacterial responses to disturbances could be especially susceptible to artifacts from exDNA pools because bacterial death can increase necromass-derived exDNA. Here, we test whether exDNA alters our ability to characterize bacterial abundance, diversity, and community structure following soil drying-rewetting, a stressful and likely lethal, pulse disturbance (12–14). We hypothesize that exDNA pools will increase after rewetting due to death, causing underestimates of bacterial mortality and misrepresentation of community shifts. To expand the generality of the study, we also test whether certain attributes of microbial communities, specifically exDNA pool size and similarity of exDNA to live communities, can predict when exDNA is likely to bias conclusions in other studies (6,7,15).

We characterized microbial dynamics in field soils at 3 time points: after a 28-day drought (∼6hrs before rewetting, t0), 1hr after a field-simulated 80mm rewetting event (t1) and 18hrs after rewetting (t2). At each time point we analyzed samples with and without excluding exDNA. We expected exDNA pools to increase in the first hour after rewetting and to alter conclusions about rewetting responses, but to decline after 18hrs, as increases in soil moisture stimulate exDNA decay (16). We also expected soil properties to alter exDNA dynamics and performed the experiment in two cropping systems to test this: conventionally-tilled corn monoculture and perennial switchgrass monoculture. We isolated the effects of exDNA by removing exDNA pools in one of two paired soil samples using the chemical propidium monoazide (PMAxx) which binds to exDNA and prevents amplification (modifying and improving the efficacy of methods in Carini et al. (6), Fig. S1). On both exDNA-included (+exDNA) and exDNA-removed (-exDNA) samples, we assessed soil bacterial abundance (16S copy number with quantitative PCR), community composition (MiSeq sequencing of 16S rRNA gene), and diversity over the drying-rewetting pulse disturbance.

## Results and Discussion

The ability to accurately characterize microbial community dynamics under disturbances is critical to understanding microbial ecology and function (17,18). We found that removal of exDNA more precisely quantified bacterial responses to a short-term disturbance (drying-rewetting), but inclusion of exDNA did not obscure results to a large extent. That is, removing exDNA sometimes altered statistical significance but rarely changed the direction of the response or general conclusions about community stability (Fig. 1a,b,c). Inconvenietly, the tendency for exDNA to mask responses was not very predictable by factors like exDNA pool size, the number of taxa unique to exDNA pools, or dissimilarity between +exDNA and -exDNA communities (Fig. 1d,e,f), and differed by time and soil type (Table S1). Overall, exDNA removal may not be necessary for most questions about bacterial responses (15), but may be especially important for detecting subtle disturbance-mediated shifts, dynamics of specific taxa, or when studies have low power.

**Fig. 1.**
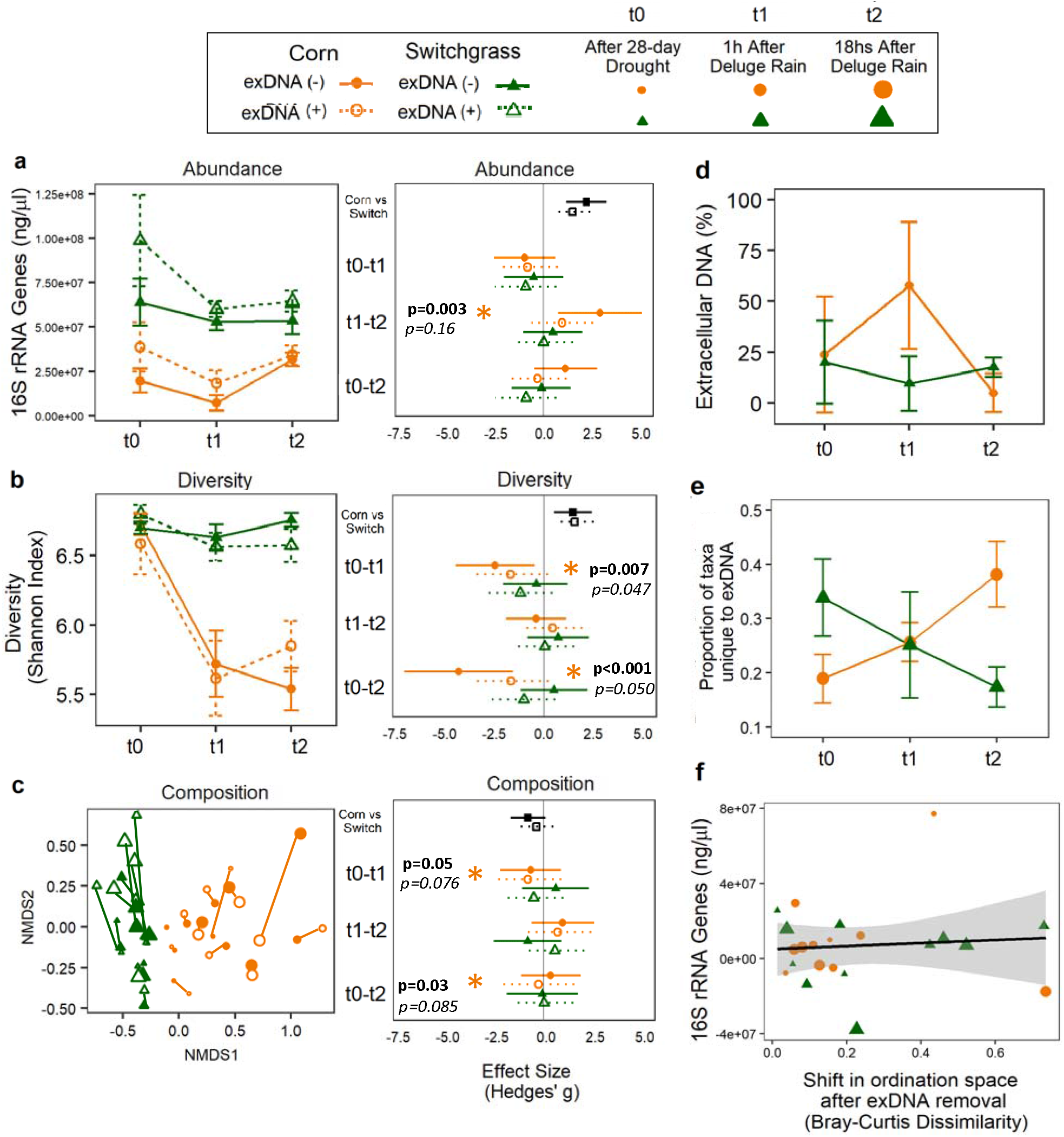
The effect of exDNA removal in corn (orange circles) and switchgrass soil (green triangles) where open points are samples with exDNA and closed points without exDNA. Bacterial responses and hedges’ g effect size for **(a)** prokaryotic abundance (number of 16S rRNA genes) **(b)** Shannon diversity and **(c)** community composition. Bacterial responses sensitive to changes in significance level (p-value cutoff) after exDNA removal are denoted by asterisks. The p-values listed on top in bold correspond to analyses that exclude exDNA and the italicized p-values listed on bottom correspond to analyses that include exDNA. **(d)** The average exDNA (%), **(e)** the proportion of OTUs unique to the exDNA pool (not present in the live community) and **(f)** the relationship between community dissimilarity and size of exDNA pool across the disturbance. Bars and points correspond to the mean, and error bars to the standard error of the mean (n=4 field replicates).

We hypothesized that exDNA would obscure changes in bacterial abundances after soil rewetting (timepoint t1) because rewetting causes bacterial death, increasing exDNA. Although bacterial populations did decline in corn soil from t0 to t1 (Fig. 1a, as measured by 16S copy number), and exDNA constituted a large perentage of the total DNA pool at this stage (Fig. 1d, 60% exDNA t1), exDNA bias was not any more prominent at this time point than in others. exDNA pools were actually larger before rewetting (t0), potentially because drought caused death as well as slowed exDNA pool degradation. Still, the slight overestimation in abundance caused by exDNA inclusion could cause some studies that included exDNA to miss the significant recovery of communities after rewetting (Fig. 1a, see p-value change t1-t2).

Inclusion of exDNA also had subtle effects on diversity and community composition (Fig. 1b,c). Removing exDNA frequently altered the composition of any one sample by a similar magnitude as commnity shifts over time (Fig. 1c, size of points show time and lines connect +exDNA and -exDNA samples), and exDNA inclusion sometimes altered significance levels for changes in composition and diversity (Fig. 1b,c, see p-value changes). However, compositional stability under drying-rewetting did not differ drastically in exDNA+ and exDNA-scenarios. In addition, certain time points were not more affected than others by including exDNA (Fig 1c, size of points), and could not be explained by the amount of exDNA (Fig. 1d), such that larger exDNA pools did not contain more unique taxa or exhbit greater dissimalrity to the live fraction of the community (Fig. 1e,f). However, analyses with only the live community highlighted certain taxa as responsive to drying-rewetting, and these were different than taxa identified in exDNA-included anlayses (10) (Table S2).

There are times when exDNA removal may be essential for accurate representation of microbial communities (6,7). While it would be helpful to predict when and where exDNA is most likely to mask responses, our study suggests this will be difficult to predict *a priori*. First, dynamics will differ across soil type (Fig. 1a,b,c). We tested two soils from the same site, but that developed different properties due to management (corn vs. switchgrass cropping systems). Soils had different moisture dynamics (Fig. S2), which likely led to distinct disturbances for microbes; drying-rewetting may have been less stressful for communities in switchgrass soil, thus there was no strong effect for exDNA to mask. Other properties likely differed as well, like pore structure, nutrient dynamics, and starting microbial communities. We do note that exDNA inclusion did not alter our ability to distinguish between corn and switchgrass microbial communities (Fig 1c, Table S1), nor did it affect our ability to distinguish between communities exposed to the drying-rewetting disturbance versus standard field conditions (Fig. S3). This suggests that exDNA is unlikely to mask large differences in microbial communities, such as those common across ecosystems or even under press disturbances (9,15).

Overall, our study suggests that exDNA removal can increase the ability to detect subtle microbial responses to pulse disturbances or changes in particular taxa, but is not likely to fundamentally change conclusions. Still, we found exDNA can alter significance levels, which would mask resistance and resilience in the many studies strictly adhering to p-value cutoffs. Finally, abiotic factors specific to the global change (e.g. temperature, moisture) and soil type are likely to affect exDNA dynamics as they alter exDNA degradation capacity and micro-scale stress responses. Although some recent evidence suggests that a significant portion of microbial diversity comes from exDNA and can alter treatments effects, our study suggests that even large compositionally distinct exDNA pools minimally bias conclusions about bacterial disturbance dynamics.

## Acknowledgements

We thank all team members of the W.K. Kellogg Great Lakes Bioenergy Research Center that helped to maintain field-scale rain out shelters. Additionally, we thank Francisca Donkor for her help conducting initial tests on the PMAxx soil treatment. We are grateful to Ashley Shade for her manuscript revisions and her recommendation to quantify effect size.

## Author Contributions

HK, KD and SE conceptualized the experimental design for soil sampling and exDNA removal. KG designed and maintained the drying rewetting experiment. HK and KD conducted the PMAxx treatment, the DNA extractions, and the qPCR. HK and SE performed the analyses and wrote the manuscript. All authors provided comments on the manuscript.

---

**Fig. S1.**
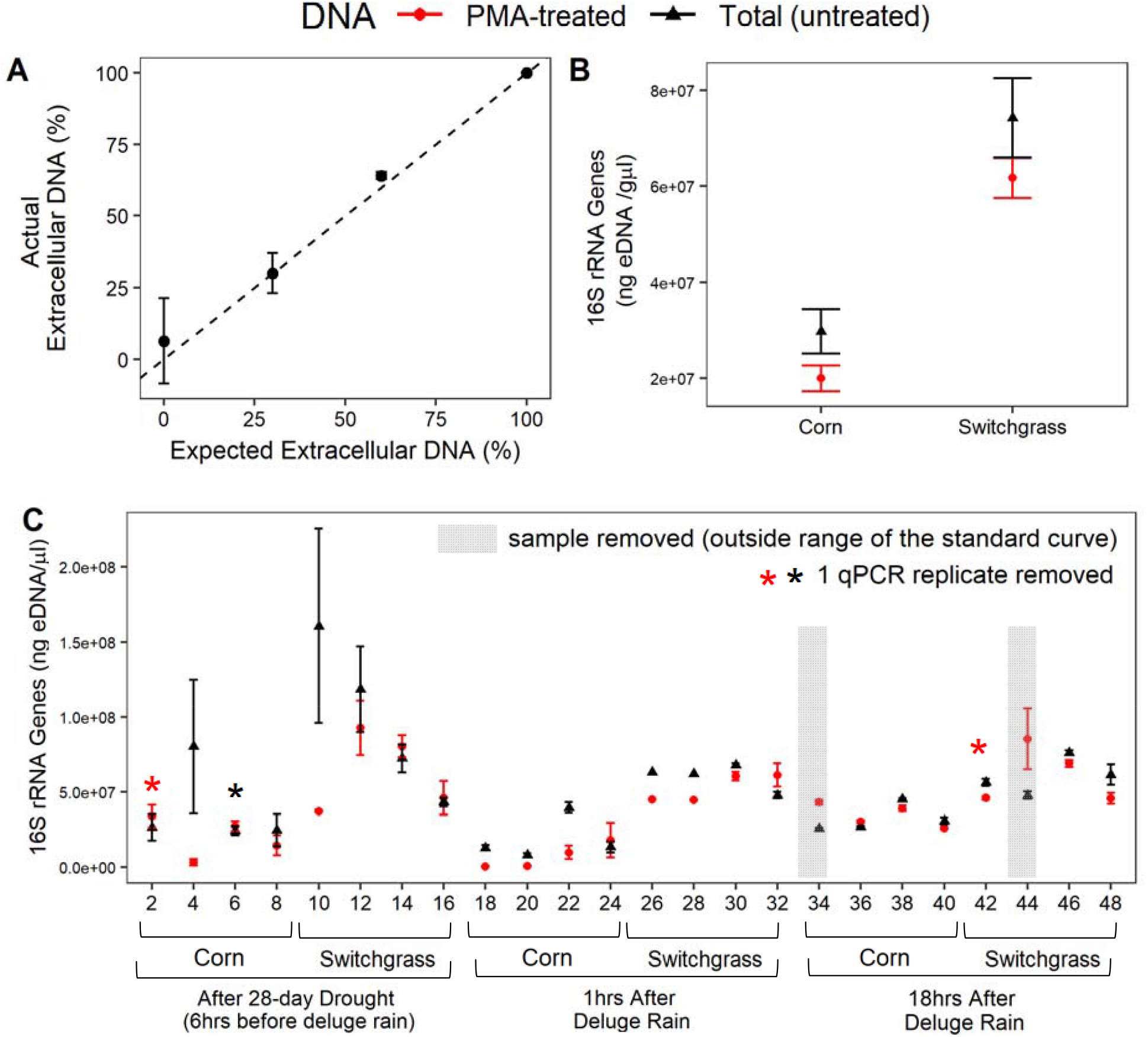
Technical aspects of exDNA removal. **A** Standard curve for the PMA-treatment in mock communities of live and dead *Pseudomonas stutzeri* inoculated into switchgrass soil (*Panicum virgatum*, variety “Cave-in-Rock”). Points show the average exDNA (%) in each mock community regressed against the expected exDNA (%), error bars show the standard error of the mean and the dashed line corresponds to x=y (n=3 soil replicates and n=3 qPCR replicates). **B** Average number of 16S rRNA gene copies in the PMA-treated (red circles) and untreated soil (black triangles) samples across the time series for corn and switchgrass soil with the error bars showing the standard error (Corn n=44 and Switchgrass n=45). **C** The average number of 16S rRNA gene copies across the 3 technical replicates (n=3 qPCR replicates) for each PMA-treated and untreated soil sample. The corresponding point in the time series and its crop type are shown below the plot. Two samples that fell outside the standard curve were removed and are highlighted in gray. Red asterisks indicate a PMA-treated technical replicate was removed and a black asterisk indicates an untreated technical replicate was removed.

**Fig. S2.**
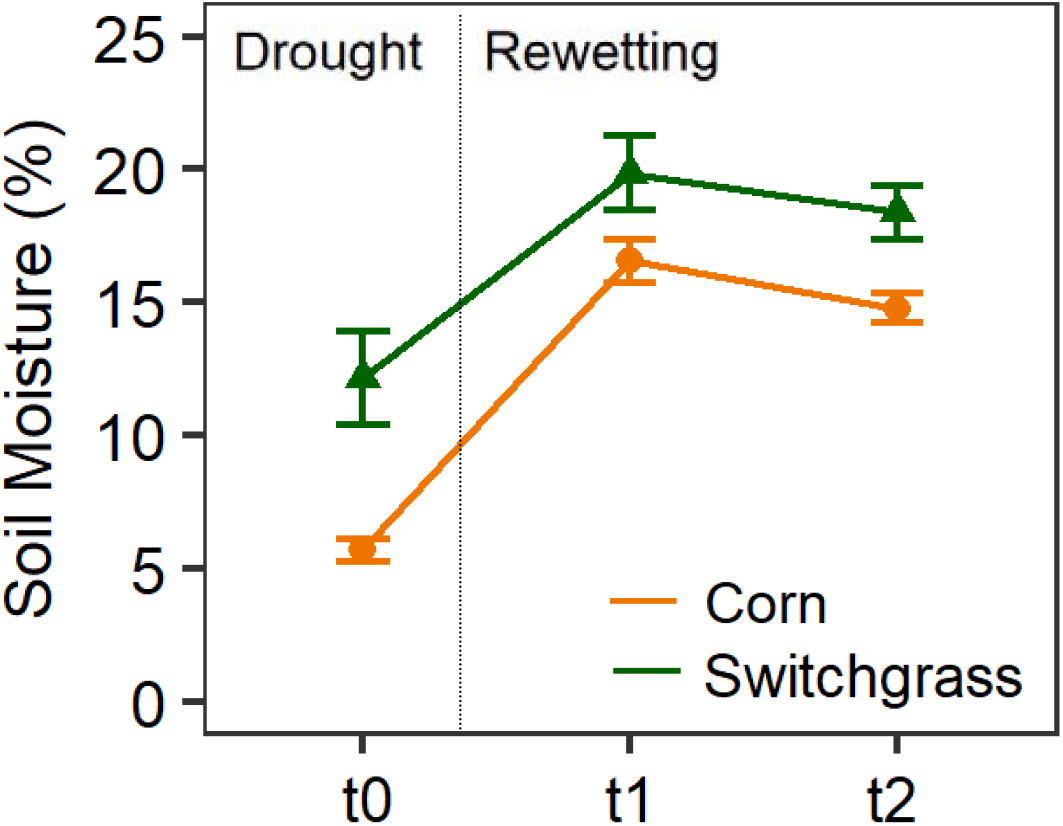
Changes in gravimetric soil moisture across the drying-rewetting disturbance for corn (orange circles) and switchgrass (green triangles) bulk soil.

**Table S1.** Expanded table of effect sizes and p-values for exDNA removal. Complete list of effect sizes and p-values for changes in abundance, diversity, and community composition across the disturbance. C corresponds to corn and S to switchgrass. (-) corresponds to values from analyses without exDNA and (+) corresponds to values from analyses that include exDNA. The rank refers to the effect of exDNA removal with 1 being the largest change in effect size after exDNA removal. P-values are only included if removing exDNA changed the significance level.

| Treatment | Measure | Time | Effect Size<br>(hedges' g) | | $\Delta$ | rank | p-value | |
| --- | --- | --- | --- | --- | --- | --- | --- | --- |
|  |  |  | (-) | (+) |  |  | (-) | (+) |
| C vs S | Composition | C vs S | -0.77 | -0.35 | 0.42 | 15 |  |  |
| C | Composition | t0-t1 | -0.64 | -0.78 | 0.14 | 19 | 0.05 | 0.076 |
| S | Composition | t0-t1 | 0.63 | -0.49 | 1.12 | 6 |  |  |
| C | Composition | t1-t2 | 0.99 | 0.74 | 0.25 | 17 |  |  |
| S | Composition | t1-t2 | -0.8 | 0.58 | 1.38 | **5** |  |  |
| C | Composition | t0-t2 | 0.38 | -0.23 | 0.61 | 13 | 0.033* | 0.085 |
| S | Composition | t0-t2 | -0.05 | 0.04 | 0.09 | 20 |  |  |
| C vs S | Abundance | C vs S | 2.23 | 1.52 | 0.71 | 11 |  |  |
| C | Abundance | t0-t1 | -0.957 | -0.81 | 0.147 | 18 |  |  |
| S | Abundance | t0-t1 | -0.488 | -0.903 | 0.415 | 16 |  |  |
| C | Abundance | t1-t2 | 2.93 | 0.991 | 1.939 | **2** | 0.003* | 0.16 |
| S | Abundance | t1-t2 | 0.499 | 0.021 | 0.478 | 14 |  |  |
| C | Abundance | t0-t2 | 1.16 | -0.29 | 1.45 | **4** |  |  |
| S | Abundance | t0-t2 | -0.09 | -0.88 | 0.79 | 9 |  |  |
| C vs S | Diversity | C vs S | 1.47 | 1.55 | 0.08 | 21 |  |  |
| C | Diversity | t0-t1 | -2.45 | -1.69 | 0.76 | 10 | 0.007 | 0.047 |
| S | Diversity | t0-t1 | -0.385 | -1.18 | 0.795 | 8 |  |  |
| C | Diversity | t1-t2 | -0.39 | 0.436 | 0.826 | 7 |  |  |
| S | Diversity | t1-t2 | 0.72 | 0.046 | 0.674 | 12 |  |  |
| C | Diversity | t0-t2 | -4.29 | -1.66 | 2.63 | **1** | 0.0004 | 0.05 |
| S | Diversity | t0-t2 | 0.51 | -1 | 1.51 | **3** |  |  |

**Table S2.** The top 15 taxa changing in response to soil rewetting. Dominant taxa across the 3 sampling points when exDNA is excluded and included in analyses. Taxa or OTUs that are incorrectly identified in the presence of exDNA are italicized.

| Taxa Responding to Soil Rewetting |  |
| --- | --- |
| (-) exDNA excluded | (+) exDNA included |
| Corn Bulk Soil |  |
| Mizugakiibacter | Mizugakiibacter |
| Sphingomonas | Thermosporotrichaceae |
| Thermosporotrichaceae | *Rhodanobacter* |
| Acidobacterium | Sphingomonas |
| Xanthomonadaceae | Acidobacterium |
| Bryobacter | Bryobacter |
| Bryobacter | *Chitinophagaceae* |
| Acidotherrmus | *Mizugakiibacter* |
| Sphingobium | *Chitinophagaceae* |
| Sphingomonas | *Castellaniella* |
| Deltaproteobacteria (GR_WP33-30) | Bryobacter |
| Xanthomonadaceae | *Acidobacteriaceae_[Subgroup_1]* |
| Acidobacteria (Subgroup_6) | *Acidobacteriaceae_[Subgroup_1]* |
| DA101_soil_group | *Acidobacteriaceae_[Subgroup_1]* |
| Sphingomonas | *Bradyrhizobium* |
| Switchgrass |  |
| DA101_soil_group | *unclassified* |
| Sphingomonas | *Bradyrhizobium* |
| Holophagae | *Luteolibacter* |
| Nitrosomonadaceae | *Sphingomonas* |
| Acidobacteria (subgroup_4) | *Flavobacterium* |
| Acidobacteria (subgroup_4) | *Cellvibrio* |
| Acidobacteria (subgroup_4) | Acidobacteria (subgroup_12) |
| Acidobacteria (subgroup_12) | *Haliangium* |
| Sphingomonas | *Variibacter* |
| Haliangium | *Pseudomonas* |
| Acidobacteria (subgroup_6) | *Chloroflexi* |
| Reyranella | *Latescibacteria* |
| DA101_soil_group | *Acidobacteria (subgroup_6)* |
| Acidobacteria (subgroup_12) | DA101_soil_group |
| Acidobacteriaceae_[Subgroup_1] | DA101_soil_group |

**Fig. S3.**
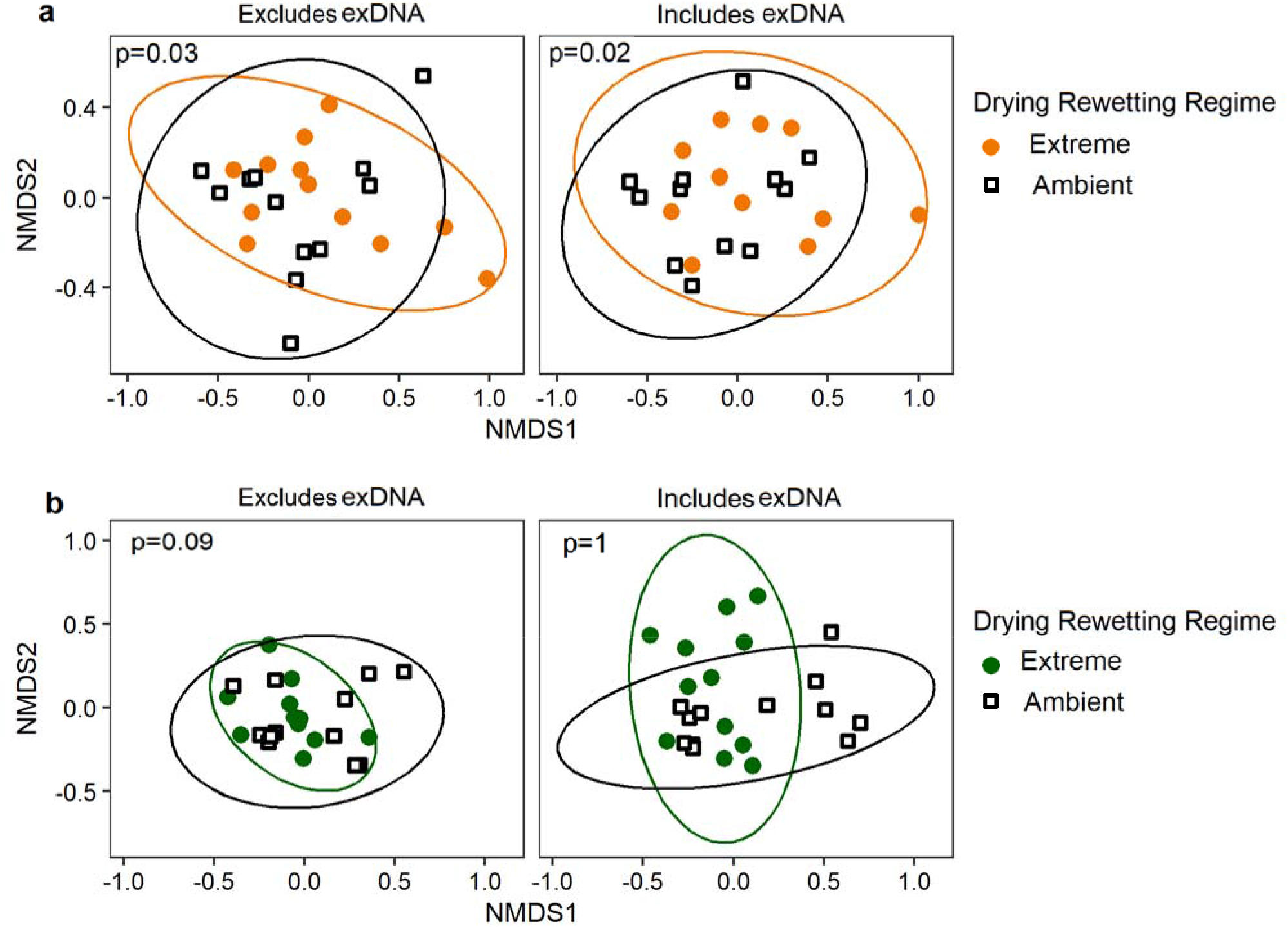
exDNA inclusion does not alter conclusions about historical exposure to ambient or extreme drying-rewetting. Community composition for **(a)** corn and **(b)** switchgrass microbial communities in plots exposed to either ambient (∼6.6mm rain every 3 days) or extreme (∼80mm rain every 28 days) drying-rewetting after one field season. Permutational multivariate analysis of variance (PERMANOVA) statistics for drying-rewetting regime differences are shown with and without exDNA removal.

## Materials and methods

### Site and soil collection

We exposed soil bacterial communities to standard or ambient drying-rewetting (6.6mm of rain every 3 days) and extreme drying-rewetting (80mm of rain every 28 days) between April 3^rd^ and September 17^th^ in experimental Biofuel Cropping System plots located at the U.S. Department of Energy’s Great Lakes Bioenergy Research Center (GLBRC) at the Kellogg Biological Station in Southwest, Michigan (42°23′47″N, 85°22′26″W, 288 m a.s.l.). On September 20^th^ of 2017 we collected soil cores (10cm depth by 5cm diameter) from continuous corn (*Zea mays*) and continuous switchgrass (*Panicum virgatum*, variety “Cave-in-Rock”) plots at three-time points; 6-hrs before, 1-hr after, and 18-hrs after a rewetting event, which also coincided with the end of a 28-day drought. The soils are well-drained mesic Typic Hapludalfs developed from glacial till and outwash consisting of co-mingled Kalamazoo (fine-loamy, mixed, semiactive) and Oshtemo (coarse-loamy, mixed, active) series with intermixed loess (1). Bulk soil was brought back to the lab, sieved at <2mm and homogenized. Gravimetric soil moisture was determined on sieved soils dried at 60°C for 72 hrs. Soil moisture increased in response to the extreme rewetting event (Fig. S2: Corn p<0.0001; Switchgrass p=0.035), while ambient rewetting had no effect on soil moisture (data not shown).

### Removal of extracellular DNA

Methods for quantifying exDNA were modified from Carini et al. (2). Two DNA extractions were completed for each soil sample using the DNeasy PowerSoil kit from Qiagen (at the time PowerSoil DNA Isolation Kit by MO BIO). A paired sample design was used where one DNA extraction tube was treated with the chemical propidium monoazide (PMAxx Dye, 20mM in H2O from Biotium) which prevents amplification of exDNA. PMAxx binds to DNA but cannot penetrate the cell wall allowing for effective removal of any DNA that is outside intact cell walls. The second DNA extraction was left untreated to quantify the total DNA pool. Both tubes were treated identically through the PMAxx activation steps. For each soil sample, 0.50 grams of soil was weighed out and evenly divided between the two DNA extraction tubes (0.25 g each). Prior to loading soil into the DNA extraction tubes, we removed the bead beating beads and the Powerbead solution that comes in the tubes. The beads were removed for the steps involving light activation of PMAxx to prevent potential cell lysis. After soil was loaded into the tube 500µl of the Powerbead solution was returned to the tube. In a dark room, 3µl of PMAxx was added to the 500µl of PowerBead Solution in the transparent bead beating tubes. Samples were homogenized for 1 minute and exposed to a 1000 W halogen light source for ten minutes while undergoing frequent homogenization (1000-Watt Halogen Telescoping Twin Head Tripod Work Light in a tinfoil lined cabinet). After light exposure, we returned the Powerbeads and the remaining PowerBead solution to the DNA extraction tube. The DNA was then extracted according to manufacturer’s instructions, however, samples treated with PMAxx were handled in low light conditions – minimal ambient light from windows – to minimize the binding of PMAxx to DNA released during the DNA extraction. The concentration of PMAxx was selected for this experiment because it optimized exDNA removal at higher concentrations of exDNA. However, this also led to increased variation in estimates of exDNA in smaller exDNA pools (Fig. S1a). In addition, previous work has shown that agricultural soils have low exDNA concentrations compared to deciduous or coniferous forest soils (Carini et al. 2016). Indicating future studies should optimize the PMAxx concentration to their specific study. However, in general, we recommend extracting DNA from PMAxx-treated samples in low light conditions or a dark room, as this reduces light-activated binding of residual PMAxx following cell lysis during DNA extraction.

### Quantitative PCR

We performed quantitative PCR in triplicate using 96-well plates on an Thermofischer thermocycler. We used 3 technical replicates for each PMA-treated and untreated soil sample (Fig. S1c). The total reaction volume was 20 µl with the following reaction mixture: 1 µl of each F and R primer (515f/806r at 10mM), 10 µl of iQ SYBR Green 2X Supermix (BIO-RAD), 4 µl sterile water, and 4 µl template DNA. The cycling conditions were: 95 °C for 15 min, followed by 40 cycles of 95°C 30 s; 50 °C 30 s; 72 °C 30 s. We generated melting curves for each run to verify product specificity by increasing the temperature from 60-95°C. Reactions were compared to standard curves developed using purified *Pseudomonas stutzeri 28a24* genomic DNA. For all qPCR reactions the linear relationship between the log of the copy number and the threshold cycle value was reported at R^2^ > 0.99. Outlier analysis was performed on qPCR replicates. We removed qPCR replicates that were 2 cycles above or below the other two replicates – this resulted in the removal of 1 qPCR replicate from 3 samples (samples 2_PMA, 6_Total and 42_PMA denoted by * in Fig. S1c). Samples 26 and 28 only had one qPCR replicate due to sample loss. Two soil samples with negative exDNA estimates were removed from the third time point analysis in both corn and switchgrass soil as they fell outside of the standard curve and were biologically inaccurate (sample 34 and 44, highlighted with gray bars in Fig. S1c). Across the entire data set, soil samples treated with PMAxx had lower variation across qPCR replicates than untreated soil samples (Fig S1b).

### Amplicon sequencing and bioinformatics

We characterized bacterial communities using high throughput barcoded sequencing on the Illumina MiSeq platform at the Research Technology Support Facility (RTSF) Genomics Core at Michigan State University. Sequencing was done in a 2 × 250bp paired end format using a MiSeq v2 500 cycle reagent cartridge. The V4 hypervariable region of the 16S rRNA gene was amplified using dual-index, Illumina compatible primers 515F and 806R as described in Kozich et al. (3). Completed libraries were normalized using Invitrogen SequalPrep DNA Normalization plates, then pooled and cleaned up using AmpureXP magnetic beads. The 16S reads were quality filtered and merged using the USEARCH pipeline (http://drive5.come/usearch/) and Cutadapt was used to remove primers and adapter bases before reads were filtered and truncated to 250bp. OTUs were clustered at the 97% sequence identity level and classified using UPARSE. Two samples with low read number were removed, making the lowest read number 7723. Singletons, chloroplasts, and mitochondria were removed (312 total) resulting in 15,062 bacterial and archaeal OTUs and 2,259,918 reads across all samples. All analyses were performed and visualized using bacterial counts that did not undergo rarefaction (4).

### Statistical analyses

We tested for differences among univariate response variables (e.g. soil moisture, Shannon diversity, and bacterial abundance or number of 16S rRNA gene copies) using Type III sum of squares ANOVA with a post-hoc Tukey HSD correction test (p < 0.05) using the lme4 and agricolae packages in R. For all analyses, we report average values across the four field replicates, except for the qPCR data, where we first calculated the number of 16S rRNA gene copies for each technical replicate (n=3 qPCR replicates, Fig. S2c). We calculated the percent exDNA by subtracting the number of 16S rRNA gene copies in the live fraction from the total number of copies, and then dividing by the total number of copies [(Total-Live)/(Total)*100].

We compared statistical differences in bacterial community composition using permutational analysis of variance (PerMANOVA) (9999 permutations) on weighted Unifrac distance matrices (Bray-Curtis distance) using the Adonis function in the vegan R package. We visualized community composition using Non-metric multidimensional scaling (NMDS) using phyloseq and ggplot2. We examined the proportion of taxa present in both the live community and exDNA pool using the Venn Diagram function in R. To assess shared membership in the exDNA pool we performed a Venn Diagram analysis on each field replicate and calculated the mean proportion of taxa specific to the exDNA pool at each time point. We report the proportion of taxa in this analysis instead of the raw number of taxa to account for changes in the total number of taxa across the disturbance. To estimate the effect of exDNA removal, we calculated standardized effect sizes on soil samples with and without exDNA, according to biascorrected Hedges’ g values (5) using the effsize package in R. We calculated the change in effect size after removing exDNA by taking the absolute value of samples with exDNA excluded minus exDNA included (as listed in Table S1).

